# An ecological explanation for hyperallometric scaling of reproduction

**DOI:** 10.1101/2021.03.12.435090

**Authors:** Tomos Potter, Anja Felmy

## Abstract

In wild populations, large individuals have disproportionately higher reproductive output than smaller individuals. Some theoretical models explain this pattern – termed reproductive hyperallometry – by individuals allocating a greater fraction of available energy towards reproductive effort as they grow. Here, we propose an ecological explanation for this observation: differences between individuals in rates of resource assimilation, where greater assimilation causes both increased reproduction and body size, resulting in reproductive hyperallometry at the level of the population. We illustrate this effect by determining the relationship between size and reproduction in wild and lab-reared Trinidadian guppies. We show that (i) reproduction increased disproportionately with body size in the wild but not in the lab, where resource competition was eliminated and food availability restricted; (ii) in the wild, hyperallometry was greatest during the wet season, when resource competition is strongest; and (iii) detection of hyperallometric scaling of reproduction at the population level was inevitable if individual differences in assimilation were ignored. We propose that ecologically-driven variation in assimilation – caused by size-dependent resource competition, niche expansion, and chance – contributes substantially to hyperallometric scaling of reproduction in natural populations. We recommend that mechanistic models incorporate such ecologically-caused variation when seeking to explain reproductive hyperallometry.

## Introduction

Across animal taxa, larger individuals reproduce more than smaller individuals of the same species (Honěk, 1993; Kingsolver & Huey, 2008; Lim et al., 2014; Ronget et al., 2018; Visman et al., 1996). While many theoretical models assume that reproduction is proportionate to body size (e.g. Kooijman, 2010; Lester et al., 2004; Roff, 1993; West et al., 2001), recent work in marine fishes has shown that larger individuals have disproportionately higher reproductive output: of 342 species, 95% display hyperallometric scaling of reproduction (Barneche et al., 2018). Reproductive hyperallometry – where large individuals produce more offspring per unit body weight than small individuals – appears to be the norm for free-living marine fishes, with potentially profound implications for the management of global fish stocks (Marshall et al., 2021). Furthermore, reproductive hyperallometry has been reported in many other taxa (e.g. freshwater fish (King, 1998), crustaceans and marine invertebrates (Thompson, 1979), true bugs (Valle et al., 1987) turtles (Mueller et al., 1998) and frogs (Gibbons & McCarthy, 1986); reviewed in Marshall & White (2019)). These widespread observations beg the question: what processes – absent from theoretical models that assume proportionate scaling of reproduction – generate these patterns?

We argue that processes occurring at (at least) two different levels of biological organization – the individual, and the population – can plausibly explain patterns of hyperallometric reproduction in natural populations. Mechanisms at both levels can be illustrated by a simple bioenergetic model of three processes central to any organism: assimilation of energy from the environment, and use of energy for either growth and somatic maintenance, or for reproduction (Fig. 1). Here we describe two non-exclusive versions of this model which result in hyperallometry of reproduction at (i) the level of the individual, and (ii) the level of the population. Although more complete individual-based bioenergetic models exist, (such as those from dynamic energy budget theory (Kooijman, 2010), or the equal fitness paradigm (Brown et al., 2018; Burger et al., 2021)), we chose to present highly simplified models to emphasize how processes operating at different levels of biological organization can generate reproductive hyperallometry.

**Figure 1:**
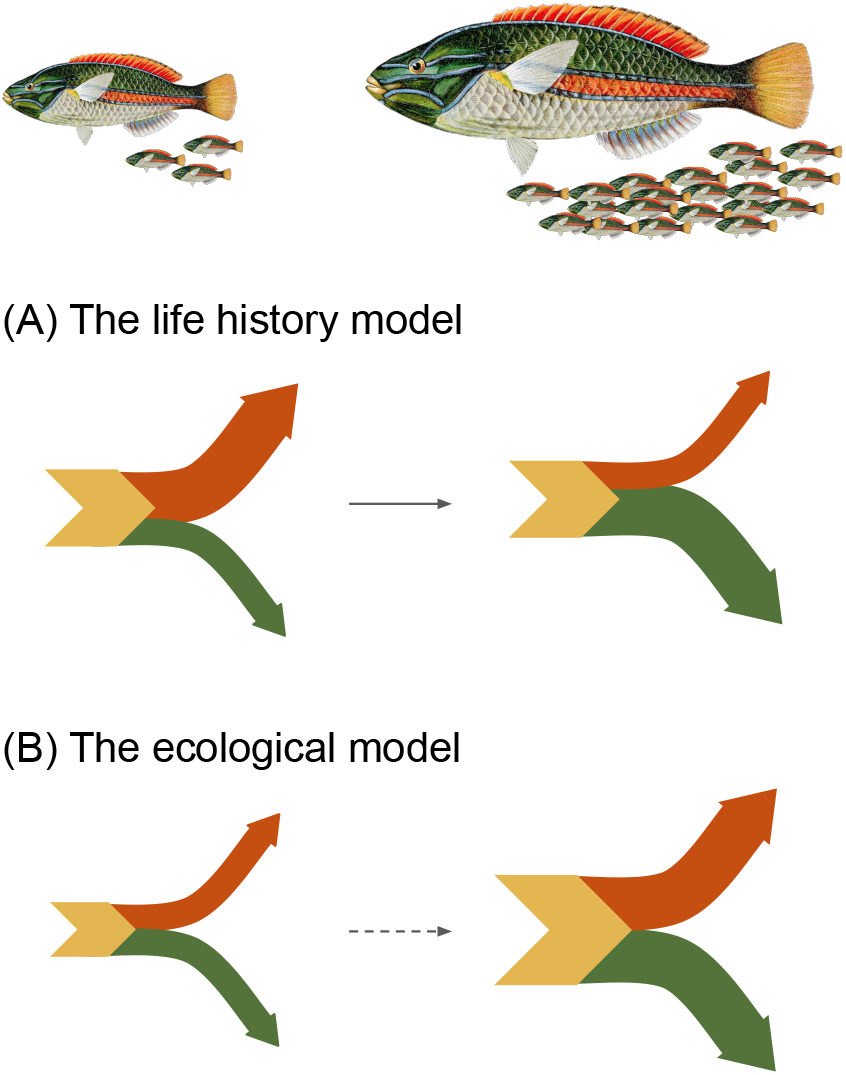
Two versions of a simple bioenergetic model of an organism which both predict hyperallometric scaling of reproduction. The organism is characterized by three energetic fluxes: assimilation of energy into the organism from the environment (yellow arrow), and use of energy for growth and somatic maintenance (green arrow) and for reproduction (red arrow). The width of arrows depicts the magnitude of fluxes. In the life history model (A) allocation to reproduction increases as the organism grows (solid black arrow). Assimilation increases with body size in absolute, but not relative terms: Marshall and White (2019) assume a sizescaling exponent for assimilation of 0.81, i.e. assimilation ~ size^0.81^. In the ecological model (B) allocation to reproduction does not change with size, but individuals with high rates of assimilation can allocate more energy to both growth and reproduction than individuals with low rates of assimilation. Although assimilation rates may increase as individuals grow (dotted arrow), individual variation in assimilation rates is sufficient for hyperallometric scaling of reproduction to arise at the population level. The fish illustration (*Stetholojulis albovittata*) is in the public domain, accessed through the Biodiversity Heritage Library.

At the level of the individual, several theoretical models predict or assume reproductive hyperallometry (Day & Taylor, 1997; Gadgil & Bossert, 1970; Kozlowski, 1996; Kozłowski & Uchmanski, 1987; Marshall & White, 2019; Quince et al., 2008). Regardless of differences in their formulations, the essence of these models is that individuals can allocate an increasing fraction of energy towards reproduction as they age (Fig. 1A). This increasing investment in reproduction can explain why larger, older individuals produce disproportionately more offspring. Predictions using this general framework match empirical observations of both reproductive hyperallometry and individual growth curves (Marshall & White, 2019). Because this mechanism depends on ontogeny, we term this the “life history model”.

In the second model we assume that the fraction of energy allocated to reproduction is fixed within an individual, but rates of assimilation vary among individuals (Fig. 1B). Individuals with higher rates of assimilation attain both larger sizes and higher reproductive output than those with lower assimilation rates. Under this model, reproductive hyperallometry emerges at the level of the population due to individual differences in assimilation rates, even if the underlying individual model assumes isometric scaling of reproduction (e.g. no ontogenetic change in energetic allocation). This is essentially the model proposed by van Noordwijk and de Jong (1986) to explain why the detection of life history trade-offs often fails at the level of individuals within populations: positive correlations between life history traits (such as reproduction and somatic growth) will occur at the population level when variance in resource assimilation affects both traits within individuals. We term this the “ecological model”, because we propose that variance in assimilation rates – and therefore the degree of reproductive hyperallometry – is frequently driven by the strength of ecological interactions.

Ecological interactions that will influence assimilation rates include resource competition, niche expansion, and the risk of predation while foraging for food. A simplifying assumption of the life history model – that individuals (of different sizes) experience the same environmental conditions (Marshall & White, 2019) – is unlikely to hold true in natural systems with respect to a key environmental variable: food availability. In wild populations, individuals are rarely equal in their ability to accrue resources, and such differences can drive divergence in realized life histories, generate population dynamics, and determine the strength of ecological interactions (Bassar et al., 2016; Coulson, 2020; Griffiths et al., 2020; Laskowski et al., 2021).

Evidence for intraspecific competition for resources is widespread among fishes, and larger body size is typically associated with stronger competitive ability (Ward et al., 2006, and references therein). Individuals may become large because their local habitat is rich in food, or because they have certain behavioral or metabolic traits that increase their rates of energy assimilation in a given environment (Auer et al., 2020; McCarthy et al., 1992, 1994; Toscano et al., 2016; Ward et al., 2004). Furthermore, once an individual becomes large, it is likely to experience further size-dependent competitive advantages over smaller individuals, under scenarios of both exploitative and interference competition (Griffiths et al., 2020; Mittelbach, 1981; Post et al., 1999; Potter et al., 2019; Ward & Krause, 2001). In addition, most fish are gape-limited feeders, which has two main consequences for size-dependent resource availability: larger fish can consume a broader range of food items (i.e. size-dependent niche expansion, Scharf et al., 2000), and they are at lower risk of predation, meaning that they can forage in areas too risky for smaller individuals (Byström et al., 2004; Magnhagen & Borcherding, 2008; Werner & Hall, 1988). As such, big fish inhabit a very different environment – in terms of food availability, past and present – than small fish within the same population.

In wild populations, disentangling the causes and effects of resource assimilation on size and reproduction is extremely challenging. Observational data on size and fecundity alone cannot, for example, directly distinguish (i) whether larger individuals allocate a greater fraction of available energy to reproduction (life history based individual-level mechanism, Fig 1A), or (ii) whether they are larger and have greater reproductive output because they have consumed more food than smaller individuals (ecological population-level mechanism, Fig 1B). By contrast, in the lab, ecological effects such as resource competition and niche expansion can be eliminated, allowing direct assessment of these questions.

Here, we assess factors that determine how reproductive output scales with body size in the Trinidadian guppy (*Poecilia reticulata*). Guppies are small, live-bearing freshwater fish common throughout streams and rivers in Trinidad and Venezuela (Magurran, 2005). Resource competition is asymmetric in guppies, with larger fish being stronger competitors (Bassar et al., 2016; Griffiths et al., 2020; Potter et al., 2019). We compared reproductive scaling exponents from observations of wild populations with those of experimental populations in the absence of the effects of resource competition and niche expansion. Specifically, we assessed how reproduction scaled with female body size (i) for different measures of reproductive output both in the wild and lab, (ii) for wild fish sampled either in the dry or (resource-poor) wet season, and (iii) for lab-reared fish fed either a high-quantity or low-quantity diet. During the wet season in Trinidad, increased water flow and flooding scours the streams of the invertebrates, algae, and detritus that guppies feed on, leading to more intense competition for resources (Auer et al., 2012; Kohler et al., 2012; Zandonà et al., 2015). In the laboratory, fish were housed individually and fed with quantified rations of food equally accessible to all size classes of fish, and so any effects of food level are independent of competitive ability. If larger guppies have disproportionately higher reproduction in both seasons (in the wild) and both food treatments (in the lab), this would support the life history model, i.e. increasing allocation to reproduction with increasing body size. By contrast, if scaling is hyperallometric in the wild, but not in the lab (in the absence of effects of competition), this would support an ecological explanation for hyperallometric scaling of reproductive output in natural populations of guppies.

## Methods

### Data

The wild data are from an observational study on variation in life histories among natural populations of guppies (Reznick, 1989, see for methodological details), supplemented with additional, previously unpublished data. Briefly, wild guppies were collected from 19 populations in the Northern Range mountains of Trinidad between 1981 and 1987 (1986 and 1987 for the previously unpublished data). Sampling took place in both the dry season (1981, 1984-1987) and wet season (1981-1983, 1985), with most populations sampled in several years (median = 3, range = 1-7, Table S1). Female guppies were dissected and weighed, and developing embryos were counted and their dry and lean dry weight recorded. In total, the wild dataset included observations of 2547 females, of which 1483 contained embryos (Table S1).

The lab data are from an experimental study of rates of senescence in female guppies (Reznick et al., 2004; 2005). In summary, guppies were collected from four natural populations and reared for two generations under lab conditions to control for differences in maternal environments. Focal females were housed in individual tanks from 30 days of age (mean = 29.6, sd = 3.9, range = 20-40) and fed quantified rations of brine shrimp nauplii or liver paste. Guppies were assigned to either a high or low food treatment. In both treatments, ration size gradually increased over the first 180 days, and was held constant thereafter. This feeding regime was designed to allow growth patterns and maximum sizes typical of wild individuals living at low or high population densities (high or low food treatments, respectively). Juvenile females were paired with a mature male to ensure that they mated as soon as they attained maturity. At each birth, females were weighed, the number, dry weight and lean dry weight of neonates was recorded, and females were re-mated. We calculated the mean somatic growth rate during pregnancy as the change in female mass between births, divided by the length of the inter-birth interval (days). Data were collected over females’ entire reproductive lifespan (mean age at final birth = 722 days, sd = 237 days). The lab dataset included 4629 observations of 211 females (Table S2).

We calculated three measures of reproductive output: fecundity (number of offspring per litter), offspring mass (total dry weight of offspring in a litter, mg), and offspring energy (total energy content of litter, J). We calculated offspring energy by multiplying the total litter protein mass (lean dry weight) and fat mass (dry weight minus lean dry weight) by appropriate energy densities (protein = 17 kJ g^−1^, fat = 37 kJ g^−1^). Additionally, for the lab data only, we could convert measures of reproductive output into reproductive powers (i.e. output per unit time) by dividing the measures by the inter-birth interval (days).

### Models

We used mixed-effects models to address five questions: (1) How do different reproductive measures scale with body size in the wild and lab, and do environmental differences (seasonality in wild populations, food level in the lab) change this relationship? (2) What factors influence the inter-birth interval in lab-reared guppies? (3) Does probability of pregnancy in wild guppies depend on size and season? (4) Do large guppies allocate relatively less energy to growth than small guppies? (5) Do estimates of reproductive scaling change if differences among individuals in assimilation rates are ignored?

Details of all models are provided in Table 1. For questions 1, 4 and 5, we modelled reproduction (*R*) as a power function of size (*z*), i.e. *R* = *az*^*β*_1_^, where *α* and *β*_1_ are the intercept and slope or scaling exponent, respectively. Reproduction is hyperallometric when *β*_1_ > 1, isometric (proportionate to body size) when *β*_1_ = 1, and hypoallometric (smaller individuals reproduce more per unit body weight) when *β*_1_ < 1. We estimated the scaling exponent by modelling the natural log of reproduction, and the main fixed effect was the natural log of body mass.

**Table 1:**
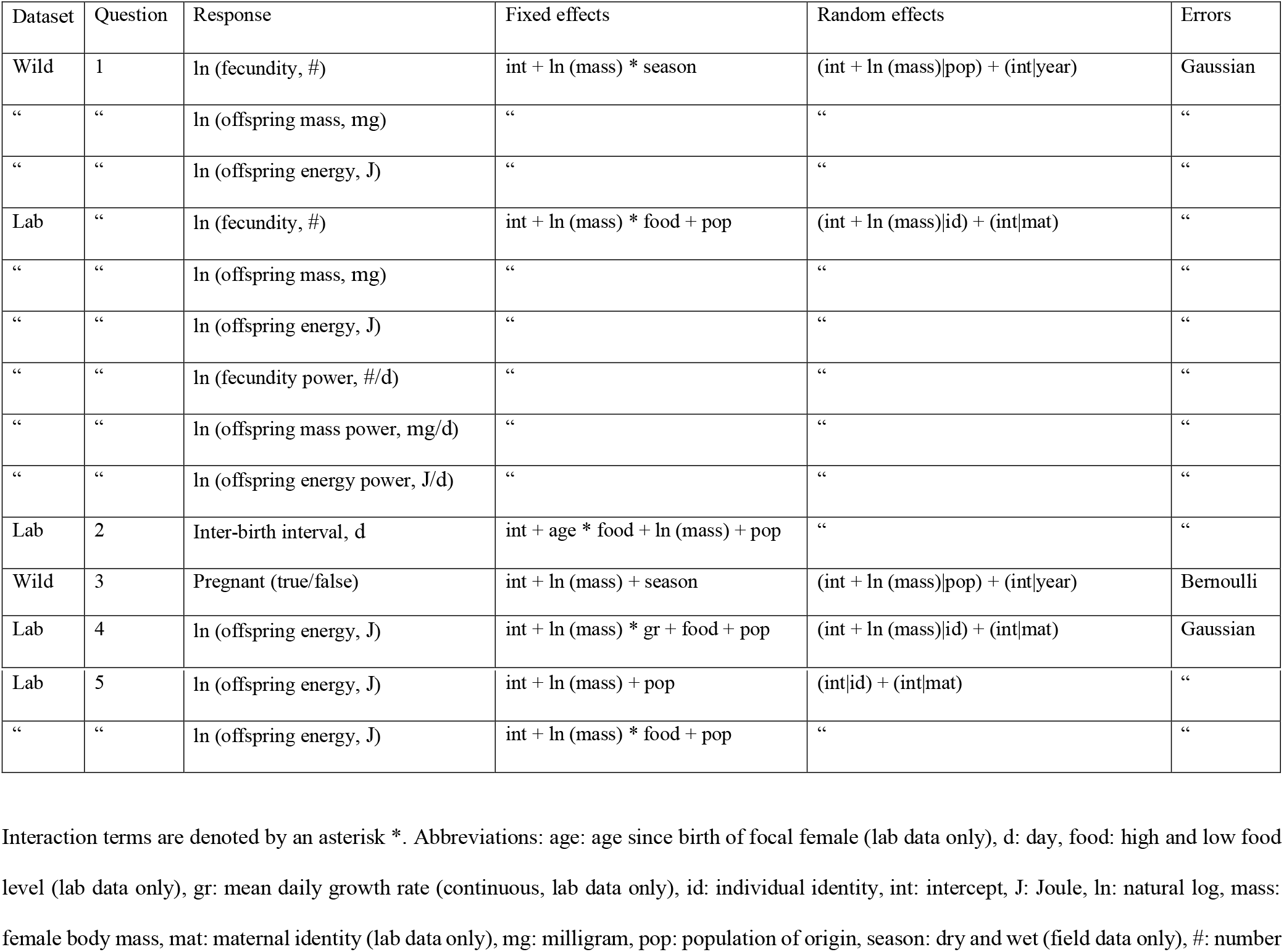
Mixed-effects models used to estimate reproductive scaling with female body size in wild and lab-reared Trinidadian guppies.

| Dataset | Question | Response | Fixed effects | Random effects | Errors |
| --- | --- | --- | --- | --- | --- |
| Wild | 1 | ln (fecundity, #) | int + ln (mass) * season | (int + ln (mass)\|pop) + (int\|year) | Gaussian |
| “ | “ | ln (offspring mass, mg) | “ | “ | “ |
| “ | “ | ln (offspring energy, J) | “ | “ | “ |
| Lab | “ | ln (fecundity, #) | int + ln (mass) * food + pop | (int + ln (mass)\|id) + (int\|mat) | “ |
| “ | “ | ln (offspring mass, mg) | “ | “ | “ |
| “ | “ | ln (offspring energy, J) | “ | “ | “ |
| “ | “ | ln (fecundity power, #/d) | “ | “ | “ |
| “ | “ | ln (offspring mass power, mg/d) | “ | “ | “ |
| “ | “ | ln (offspring energy power, J/d) | “ | “ | “ |
| Lab | 2 | Inter-birth interval, d | int + age * food + ln (mass) + pop | “ | “ |
| Wild | 3 | Pregnant (true/false) | int + ln (mass) + season | (int + ln (mass)\|pop) + (int\|year) | Bernoulli |
| Lab | 4 | ln (offspring energy, J) | int + ln (mass) * gr + food + pop | (int + ln (mass)\|id) + (int\|mat) | Gaussian |
| Lab | 5 | ln (offspring energy, J) | int + ln (mass) + pop | (int\|id) + (int\|mat) | “ |
| “ | “ | ln (offspring energy, J) | int + ln (mass) * food + pop | “ | “ |

Interaction terms are denoted by an asterisk \*. Abbreviations: age: age since birth of focal female (lab data only), d: day, food: high and low food level (lab data only), gr: mean daily growth rate (continuous, lab data only), id: individual identity, int: intercept, J: Joule, ln: natural log, mass: female body mass, mat: maternal identity (lab data only), mg: milligram, pop: population of origin, season: dry and wet (field data only), #: number

Question 1 was modelled using all reproductive measures available for lab (6) and wild data (3). To test the relationship between environmental factors and the scaling exponent *β*_1_, we included either season (wild data) or food treatment (lab data) as fixed effects, and their interactions with ln(mass).

Questions 2 and 3 concern factors influencing components of reproduction other than output: the interval between successive births in the lab, and the probability of being pregnant when sampled in the wild, respectively (details in Table 1 and Supporting Methods).

For questions 4 and 5, we modelled total offspring energy content per litter, because this reproductive measure was available in both lab and wild data and is equivalent to that used by Barneche *et al.* (2018). To assess whether energy allocation to growth decreases with increasing size in the lab (question 4), we included an interaction between somatic growth rate and ln(mass) (Table 1). Under the life history model, reproduction and growth should correlate negatively, and this relationship should be strongest at larger body sizes. This would correspond to a negative main effect of growth and a negative interaction term of growth with body size. To address question 5, we modelled reproduction of a cohort of guppies in the lab (aged 200-300 days) as a power function of size, both with and without accounting for food treatments (Table 1).

All analyses were performed using R version 3.5.1 (R Core Team, 2019). We used appropriate random effects structures to account for non-independence of data (Table 1 and Supporting Methods). We fit our models in a Bayesian framework, using the R package brms version 2.14.4 (Bürkner, 2018), which makes use of the probabilistic programming language Stan (Carpenter et al., 2017). Details of priors and modelling routines are given in the Supporting Methods.

Figures 2, 3 and S1 were produced using R packages ggplot2 (Wickham, 2011), ggridges (Wilke, 2018) and patchwork (Pederson, 2020).

**Figure 2:**
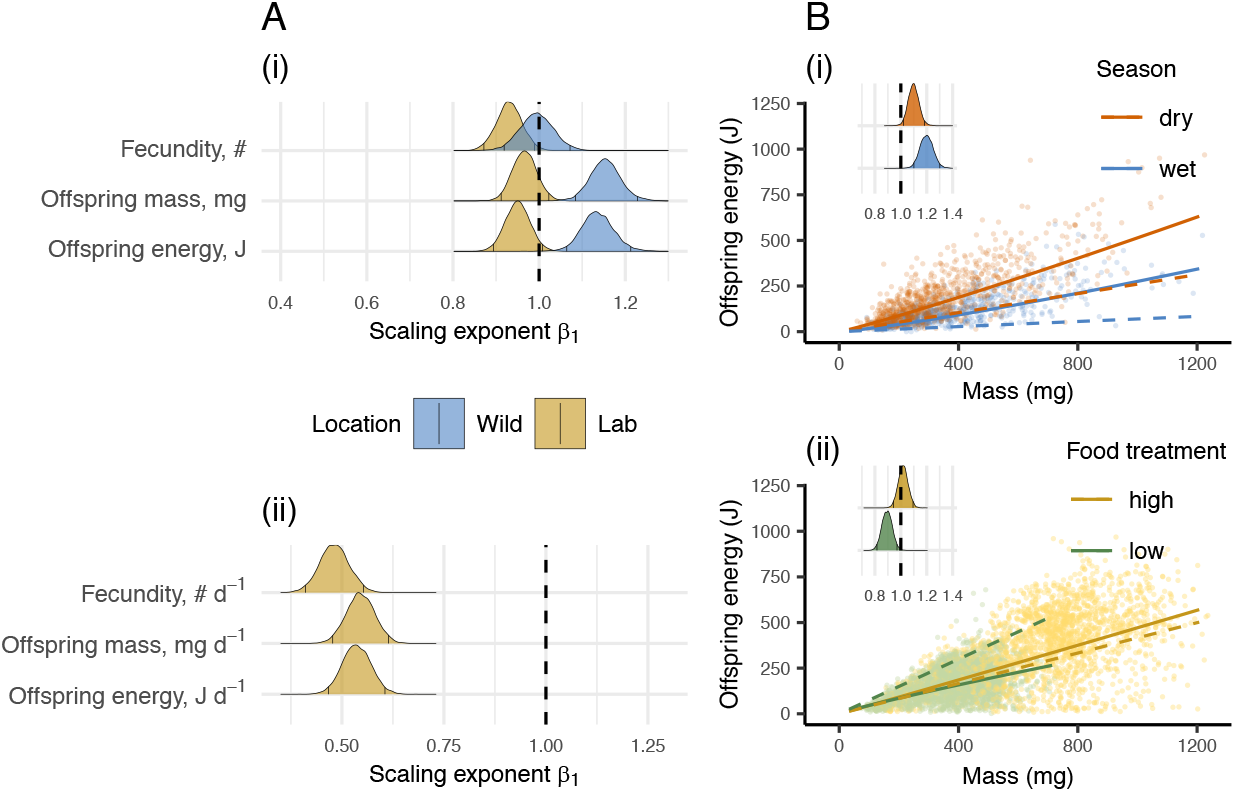
The scaling of reproduction with female body size differs between wild and lab-reared populations of Trinidadian guppies, reproductive measure, and environmental conditions. (A) shows the posterior distributions of scaling exponents using three reproductive measures, expressed (i) per litter, and (ii) per unit time (only available for lab data). Vertical black lines denote 95% credible intervals. Posteriors are given for wild (blue) and lab (yellow) populations, and are averaged across seasons (wild) and food treatments (lab), weighted by the number of observations in each season/food treatment. (B) shows offspring energy content per litter by body size in (i) wild guppies in the dry (red) and wet season (blue) (1485 datapoints, one per individual), and in (ii) lab-reared guppies at high (yellow) or low food (green) (4629 datapoints from 211 individuals). Insets show the posterior distribution of the scaling exponents. Colored lines show predicted reproduction given the estimated scaling exponents (solid) or assuming isometric scaling of reproduction, i.e. *β*_1_= 1 (dashed).

**Figure 3:**
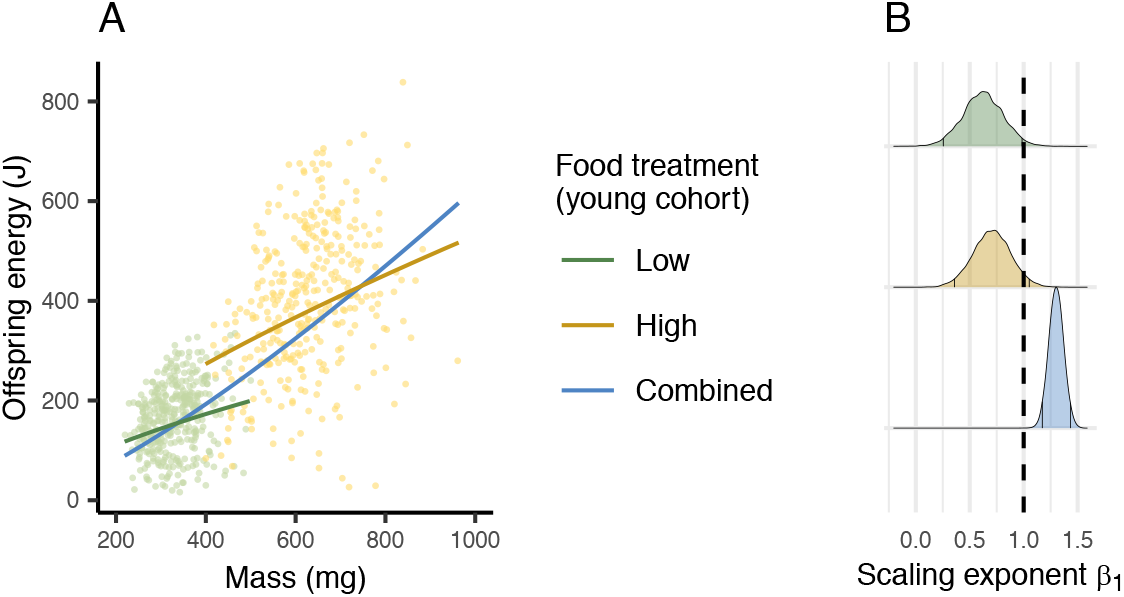
Hyperallometric scaling of reproduction was detected when differences in food availability were ignored. (A) The relationship between reproduction (offspring energy content per litter) and female body size (mass) for a cohort of lab-reared guppies aged 200 to 300 days (total N = 729), maintained at a low (green) or high (yellow) level of food availability. Lines show predicted reproduction when food levels are modelled (green and yellow) and when they are not (blue). (B) The reproductive scaling exponent *β*_1_ was greater than 1 when data from both food levels were combined (i.e. reproduction appeared hyperallometric), but exponents were less than 1 when differences in food availability were included in the model.

## Results

Parameter values reported are means of the posterior distribution, with 95% credible intervals given in brackets. Reproductive scaling is hyperallometric when the lower bound of the credible interval exceeds 1, isometric when the interval contains 1, and hypoallometric when the upper bound is less than 1.

### Environmental effects on the scaling of reproduction with body size

In the wild, mean reproductive output was hyperallometric for offspring mass (*β*_1*M*_= 1.15 [1.08; 1.23]) and offspring energy (*β*_1*E*_= 1.14 [1.06; 1.21]), but isometric for fecundity (*β*_1*F*_= 0.99 [0.92; 1.07]) (Fig. 2Ai).

In the lab, mean reproduction scaled hypoallometrically for fecundity (*β*_1*F*_= 0.93 [0.87; 0.99]), isometrically for offspring mass and energy (*β*_1*M*_= 0.97 [0.91; 1.02]; *β*_1*E*_= 0.95 [0.89; 1.01]; Fig. 2A.i), and hypoallometrically for all measures of reproductive power (*β*_1*FP*_= 0.48 [0.41; 0.55]; *β*_1*MP*_= 0.54 [0.48; 0.61]; *β*_1*EP*_= 0.54 [0.47; 0.60]; Fig. 2A.ii). This reduction in how size scales with reproductive power relative to reproductive output corresponded to a linear increase in inter-birth intervals with age: the interval was 40% longer in the oldest fish relative to the youngest (intercept = 25.4 d [23.8; 27.0], *β_age_* =0.010 [0.008; 0.013]). Interbirth intervals were not affected by female mass, the food treatment, or the interaction between age and food (*β*_*ln* (*mass*)_= −0.13 [−1.14; 0.91], *β_food low_*= 0.75 [−0.39; 1.88], *β*_*ln*(*age*):*food low*_= −0.001 [−0.004; 0.001]).

For wild populations, reproduction increased disproportionately to body size in both seasons, but hyperallometry was stronger in the wet season (offspring energy J, *β*_1 *dry*_= 1.10 [1.02; 1.18]; *β*_1 *wet*_= 1.20 [1.10; 1.30]; difference = 0.10 [0.00; 0.20], Fig. 2B.i). In the lab, reproduction scaled isometrically at high food, but hypoallometrically at low food (offspring energy J, *β*_1 *high*_= 1.02 [0.94; 1.09]; *β*_1 *low*_= 0.89 [0.82; 0.97], Fig. 2B.ii). These patterns - *β*_1 *wet*_ > *β*_1 *dry*_ in wild populations, and *β*_1 *high*_ > *β*_1 *low*_ in the lab – were consistent across all measures of reproductive output (data not shown).

### Correcting for likelihood of pregnancy intensifies hyperallometry in the wild

In the wild, pregnant females were larger on average than females without developing embryos, especially in the wet season. Consequently, reproductive hyperallometry was even stronger when accounting for size- and season-dependent pregnancy probability: the reproductive output of large females (1000 mg) was 5.2 times higher than that of small females (200 mg) in the dry season, and 10.0 times higher in the wet season (Supporting Results, Fig. S1).

### Variation in assimilation rates as a potential mechanisms underlying reproductive hyperallometry

In the lab, we found no evidence that larger fish allocate relatively more energy to reproduction over growth: when we included growth rate in our model, its main effect was negative (*β_G_*= −0.63 [−0.84; −0.44]), but its interaction with body size was positive (*β*_*G*:*z*_= 0.10 [0.07; 0.13]), meaning that growth rate and reproduction co-varied negatively in small individuals, but positively in large individuals.

Guppies reared under the high food treatment grew to larger sizes and had higher reproductive output than those in the same age cohort in the low-food treatment (Fig. 3). Although reproduction scaled isometrically at high food (*β*_1 *high*_ = 0.71 [0.36; 1.05]) and hypoallometrically at low food (*β*_1 *low*_= 0.62 [0.25; 0.98]), reproduction was hyperallometric when calculated across both food treatments (*β*_1 *both*_= 1.30 [1.17; 1.43], Fig. 3).

## Discussion

What mechanisms can explain the patterns of reproductive hyperallometry so widespread among wild populations? To date, such patterns have primarily been ascribed to an ontogenetic shift in allocation of resources within individuals (i.e. an individual-level mechanism, the life history model, Fig. 1A). We propose an alternative mechanism: individual differences in energy assimilation (i.e. a population-level mechanism, the ecological model, Fig. 1B). These two mechanisms, operating at different levels of biological organization, are not mutually exclusive. Both may contribute to observations of reproductive hyperallometry in wild populations, and the relative contribution of each mechanism is likely to depend on the study system. However, our results suggest that the ecological model better explains observations of reproductive hyperallometry in wild populations of Trinidadian guppies. Our argument is supported by three main findings. First, the strength of reproductive hyperallometry was environment-dependent (Fig. 2B.i, Fig S1). Second, in a laboratory setting devoid of resource competition and niche shifts, reproduction was not hyperallometric (Fig. 2A, Fig 2B.ii). Third, the detection of reproductive hyperallometry was inevitable when differences in energetic assimilation were ignored (Fig. 3). We expand on these points in the following sections.

### Environmental conditions affect how reproduction scales with body size

Our results add to the list of examples of reproductive hyperallometry in wild populations (Barneche et al., 2018; Marshall & White, 2019). Differences in reproductive output per unit body mass in wild guppies were driven by larger individuals producing disproportionately larger, rather than more, offspring (Fig. 2A.i), implying that total offspring biomass per litter reflects the assimilated energy by mothers. We show that the strength of hyperallometry was environment-dependent: big guppies had an even greater reproductive advantage over smaller ones during the challenging wet season (Fig. 2B.i). This result is striking, because some versions of the life history model predict that reproductive hyperallometry should be weaker during unfavorable seasons (Kozlowski, 1996; Kozłowski & Uchmanski, 1987) – under optimized energetic allocation, individuals should invest more in growth (and less in reproduction) when living conditions are challenging. Here, we argue that population-level reproductive hyperallometry increases during the wet season because resource competition is more intense (Reznick et al., 2019). When individuals are unequal competitors – as is the case in natural populations of guppies (Griffiths et al., 2020) – more intense resource competition will by definition increase variance in assimilation rates. Under the ecological model (Fig. 1B), this increase in competitive intensity explains the stronger reproductive hyperallometry in wild guppies during the wet season.

Not only did reproductive output decrease in pregnant guppies during the wet season, but smaller females were far less likely than larger females to be pregnant in the first place: when we accounted for the probability of pregnancy, the effects on reproduction of size and season increased drastically (Fig. S1B.ii). Although this measure of hyperallometry is not directly comparable with standard values of *β*_1_ (our reproduction function was non-linear on the log scale), across a typical range of adult sizes, the pregnancy-corrected estimates corresponded to values of *β*_1_ of ~5 and ~10: the reproductive output of large (1000 mg) guppies per unit body weight was five times that of a small (200 mg) guppy in the dry season, but ten times greater in the wet season. This degree of reproductive hyperallometry far exceeds those reported in other species (Barneche et al., 2018; Marshall & White, 2019), and highlights the strong effect of seasonality on the relationship between size and reproduction in wild guppies.

### No evidence for increased allocation to reproduction with increased body size

The life history model could explain the increased value of *β*_1_ in the wet season if large individuals allocate even more energy to reproduction when resources are limited. The reallocation would need to be substantial, given that per capita food availability is reduced in the wet season (Auer et al., 2012; Kohler et al., 2012). Our lab results demonstrate that this is not the case: under low-food conditions reproductive output scaled hypoallometrically, i.e. smaller fish had higher reproductive output per unit body weight, whereas reproduction was proportionate to body size at high food (Fig. 2B.ii). However, when we considered reproductive power (i.e. reproductive output per unit time), reproductive scaling was strongly hypoallometric at both food levels, as larger females gave birth less frequently (Fig. 2A.ii). This suggests that guppies became increasingly food-limited as they grew, even at high food, and compensated for the increasing costs of somatic maintenance with larger body size by diverting energy away from reproduction. This is interesting for two reasons. First, it implies that energy allocation is not fixed (as assumed in the ecological model) but rather responds to environmental conditions, although here the direction of change was opposite to that predicted by the life history model. Second, it suggests that reproductive hyperallometry in wild guppies arose precisely because large females could feed as much as they liked (see “Limitations” below).

The expression of life history traits in guppies often strongly depends on food availability (Felmy et al., 2021). Hence it was unsurprising that guppies in the high food treatment were larger and reproduced more than same-aged individuals in the low food treatment (Fig. 3A). Within food treatments, mean values of *β*_1_ were less than 1 (Fig. 3). However, when estimated across both treatments, *β*_1_ was greater than 1 (Fig. 3), showing how reproductive hyperallometry can erroneously be detected at the level of the population if individual differences in assimilation rates are not taken into account.

### Causality, competition, and alternative explanations

The change from *β*_1_ < 1 to *β*_1_ > 1 (Fig. 3), and more generally the observation of positive correlations among life history traits (van Noordwijk & de Jong, 1986), exemplify the Yule-Simpson effect: a change in a covariance conditional on a third factor (Simpson, 1951; Yule, 1903). Yule-Simpson effects are common in regression analyses, occurring when covariances are estimated over aggregated data structured by a third variable (Blyth, 1972; Clark et al., 2011; Pearl, 2014). In such cases, the covariance captures a non-causal association between the two variables of interest, which is in fact conditional on a third variable. Here, the variables of interest were size and reproduction, and the third variable was resource assimilation. The life history model (Fig. 1A) assumes a causal effect of size on reproduction: bigger individuals allocate more energy to reproduction. By contrast, the ecological model (Fig. 1B) assumes a causal effect of assimilation on both growth and reproduction.

In wild populations, the causal relationship between size and assimilation rate is more nuanced than that: a large size may increase assimilation disproportionately (e.g. through increased competitive ability and niche shifts, see Introduction), resulting in a positive feedback loop between assimilation rate and body size. In this case, size has an indirect causal relationship on reproduction, through size-dependent increases in assimilation. Furthermore, competition for finite resources is a zero-sum game: food eaten by one individual cannot be eaten by its competitors. This means that individual assimilation rates will depend on competitive ability. Here, size is key: in lab-based competition trials, large guppies suppressed the somatic growth of smaller individuals more than vice-versa (Potter et al., 2019); in the wild, individual growth and survival depended on the relative size of competitors (Griffiths et al., 2020).

While size-dependent competitive ability will lead to reproductive hyperallometry, individual differences in resource assimilation can also result from stochastic processes, environmental heterogeneity, and genetic differences among individuals leading to variation in physiology, development, and behavior. Moreover, differential trait-mediated mortality, where growth rates, reproductive success, and survival all correlate positively, can also cause reproductive hyperallometry at the level of the population. Nevertheless, given that size-structured competitive interactions are common in fish (Ward et al., 2006), the strength of resource competition is likely a key determinant of the magnitude of reproductive hyperallometry in wild populations.

### Limitations

Our study considered a single species whose small body size and viviparity set it apart from many other fishes. We thus do not know whether individual differences in assimilation are as large a contributor to reproductive hyperallometry in other species as they are in Trinidadian guppies. Neither is it clear whether reproductive scaling exponents are generally higher in the wild than in the lab. However, a recent aquaria-based study of the European sea bass (*Dicentrarchus labrax*) reported hypoallometric scaling of reproduction (*β*_1_= 0.84) under constant food availability (Sadoul et al., 2020), whereas wild populations of this species show hyperallometric scaling of reproduction (*β*_1_= 1.34, Barneche et al., 2018). This and another study have predicted that, under the framework of dynamic energy budget theory (Kooijman, 2010), differences within a population in food availability or ambient temperature can generate patterns of reproductive hyperallometry (Kearney, 2019; Sadoul et al., 2020). These empirical observations and theoretical predictions are consistent with our findings here.

Even in guppies, it remains to be seen which aspect of the laboratory environment prevented large females from reproducing hyperallometrically. By housing fish individually and keeping food rations constant from age 180 days upwards, the experiment not only eliminated resource competition but also led to food limitation in larger size classes. Future studies must explore whether reproductive hyperallometry ensues if laboratory fish are allowed to compete for food (e.g. by keeping them in groups), and/or if they are reared individually but fed *ad libitum* throughout their lives.

Another potential contributor to the observed difference in scaling exponents between lab and field populations was selection bias due to differential mortality in the wild-caught samples: while in the lab we obtained data from a random sample of females over their full lifespan, large females in the wild are rare, and likely represent a non-random subset (in terms of attributes) of the population as a whole. Indeed, only 15% of large (>800 mg) wild females (N = 66) had at least one litter with low energy content (<150 J), compared to 48% (N = 60) in the lab (Fig. 2B). This difference further emphasizes the role of ecological interactions in determining reproductive hyperallometry: it is likely that those lab-reared large guppies with low reproduction (and presumably low assimilation/digestive efficiencies) would not have attained a large size under competitive conditions.

The vast majority (98%) of data used by Barneche *et al.* (2018) were from wild populations. Without any additional information on relevant ecological and environmental conditions, we argue that one cannot infer the cause of reproductive hyperallometry from such data. Interestingly, the most extreme values of hyperallometric scaling reported by Marshall and White (2019) do not come from observations of wild populations: they come from a lab-based study on the effects of experimental “crowding” on the growth and reproduction of leafhoppers (Valle et al., 1987). This scenario – where scaling exponents were measured across multiple groups differing in the intensity of resource competition – is precisely where the ecological model (and the Yule-Simpson effect) would predict the greatest degree of reproductive hyperallometry.

### Conclusions and broader implications

Reproductive hyperallometry is ubiquitous across many taxa (Barneche et al., 2018; Marshall & White, 2019). These observations have led to recent calls to abandon theoretical models that assume proportionate scaling of reproduction with body size at the level of the individual (Marshall & White, 2019, although see also Kearney 2019). Here, we suggest that these calls may be too hasty: we demonstrate how reproductive hyperallometry at the population-level can occur – even when the underlying model at the individual level assumes proportionate scaling – through differences among individuals in assimilation rates. We do not argue that individual-level models that predict hyperallometry are wrong; instead, we illustrate how plausible population-level mechanisms can contribute to observations of reproductive hyperallometry. For a better mechanistic understanding of this phenomenon, we argue that there is an urgent need to incorporate differences in assimilation rates between individuals into life history models.

In natural populations, it is inevitable that individuals differ in their rates of resource acquisition and assimilation. Whether through trait-mediated differences or chance, two free-living individuals will never have equivalent amounts of assimilated energy to fuel their metabolism, grow, and reproduce. The effects of individual differences in resource assimilation are non-trivial. Resource-dependent expression of life history traits often exceeds the effect of evolved differences between locally adapted populations (Felmy et al., 2021). Trait-dependent differences in assimilation rates can flip predictions about life history evolution (Laskowski et al., 2021), generate eco-evolutionary dynamics (Coulson, 2020), and contribute to the evolution of coexistence (Anaya-Rojas et al., 2021). Understanding how and why individuals differ in rates of resource assimilation is central to understanding variation in demographic rates that underpin ecological and evolutionary dynamics in wild populations.

## Author contributions

TP co-conceived the study, developed and performed the analyses, and wrote the first draft of the manuscript. AF co-conceived and supervised the study, and interpreted results with TP. Both authors reviewed and edited subsequent versions of the manuscript.

## Data accessibility statement

All data associated with this manuscript will be archived at Dryad Digital Repository following manuscript acceptance.

## Acknowledgements

We are very grateful to David Reznick for providing the datasets used here, to Tim Coulson and his group for incredibly helpful discussions, and to Dustin Marshall for a lively email discussion which inspired this study. We thank three anonymous reviewers who provided valuable feedback on an earlier version of the manuscript. TP received funding from the Natural Environment Research Council (United Kingdom) through the Oxford Environmental Research Doctoral Training Program, and from a Lamb and Flag scholarship from St John’s College, Oxford.

