## Supplementary Information for "An ecological explanation for hyperallometric scaling of reproduction"

### Supporting Methods: Models

Questions 2 and 3: For question 2, we included ln(mass) and the interaction between age and food level as fixed effects in our model of the inter-birth interval. For question 3, we estimated the probability of a wild-caught individual containing zero developing embryos as a function of size and season. For this model we fit a logistic regression of the binary response (presence/absence of developing embryos), and included season and the natural log of mass as fixed effects (Table 1).

Random effects structures: For all models fit, we included random effects terms to account for non-independence of data. One obvious source is the population of origin of the guppies: natural guppy populations differ from one another genetically, and in terms of the intensity of predation risk and other ecological factors (Magurran, 2005). Because there were only 4 sampling populations in the lab data, but 19 in the wild dataset, we treated sampling population as a fixed effect in the lab models, but as a random effect in the lab models (Table 1). For wild data, we allowed the intercept and slope to vary by sampling population (int + ln(mass)|pop). For wild data, we also included a random intercept for sampling year (int|year). For lab data, we allowed the intercept and slope to vary by individual identity (int + ln(mass)|id) to account for repeated measures on individuals, and we fit a random intercept to account for shared maternal identity among siblings in the study (int|mat). To answer question 5, we used a subset of the lab data, solely consisting of data on the size and reproduction of fish aged 200-300 days. As there were fewer observations per individual, we did not fit a random slope in these models. Models addressing questions 1, 2, 4, and 5 all used a Gaussian error structure. To address question 3, we fit a logistic regression with a Bernouilli error distribution. Unlike our other analyses of the wild dataset, the logistic regression included both pregnant and non-pregnant individuals.

Priors: In the Gaussian models (Table 1, questions 1, 2, 4, and 5), for the intercept and fixed effect coefficients we specified normal priors with a mean of 0 and standard deviation of 1 (sd = 5 for the intercept). For the standard deviation of random effects and residual variance, we specified cauchy priors with means of 0 and standard deviations of 1. For the correlation of random effects, we specified an LKJ prior with a value of 1, which assumes all correlational structures are equally likely (Bürkner, 2018). For the logistic regression model we used the "get_prior" function in brms to obtain appropriate default priors (Bürkner, 2018).

Modelling routines: For each model, we ran four Monte Carlo Markov chains, with 2000 samples retained from each chain after a burn-in period of 1000 samples, yielding a combined total of 8000 samples of the posterior distribution. We confirmed that chains had converged by visually checking the traceplots, and by confirming that scale reduction factors $\hat{R} < 1:01$. Effective samples sizes for all parameters were greater than 10% of the total number of samples.

### Supporting Results: Correcting for likelihood of pregnancy in the wild

In the wild, pregnant females were larger on average than females without developing embryos: 337 mg [328;347] vs. 154 mg [141;167] in the dry season, and 420 mg [406; 436] vs. 216 mg [199;233] in the wet season (Fig. S1A). The probability of pregnancy increased with size, especially in the wet season (logistic regression, intercept = -16.07 [-18.17; -14.12], $\beta_{\ln\,(mass)}$= 3.16 [2.86; 3.50], $\beta_{wet}$= -0.87 [-1.20; -0.53], Fig. S1B.i). This meant that a small female (200 mg) had a 45% chance of being pregnant in the wet season, but a 66% chance in the dry season. Accounting for size- and season-dependent pregnancy probability changed the predicted relationship between size and reproduction in the wild: large fish had an even greater relative advantage over small fish in terms of reproduction (Fig. S1B.ii). Accounting for the probability of being pregnant, the mean reproductive output (total energy content per litter) of a 200 mg female was 19.2 J in the dry season, but only 5.3 J in the wet season; for a 1000 mg female, it was 503.9 J in the dry season, and 263.9 J in the wet season. Per unit body weight, the reproductive output of a 1000 mg female was 5.2 times higher than that of a 200 mg guppy in the dry season, and 10.0 times higher during the wet season.

### Supporting Tables

**Table S1: Sample sizes for the wild dataset, by population sampling code (19 locations) and year (7 sampling years).** Numbers given in square brackets are samples collected during the wet season. Total sample size is 2547 sets of observations, one set per individual. Data collected from 1981-1985 are from Reznick (1989). Data collected in 1986 and 1987 have not previously been published.

| Population code | Years sampled | | | | | | |
|---|---|---|---|---|---|---|---|
| | 1981 | 1982 | 1983 | 1984 | 1985 | 1986 | 1987 |
| 4 | 29 | [36] | | 33 | 32 [34] | 38 | |
| 5 | 36 [36] | | [38] | 29 | | | |
| 8 | 15 [27] | | [25] | 43 | | 34 | |
| 10 | 12 [42] | | [30] | 39 | 37 [36] | | 45 |
| 15 | 11 [26] | | [35] | 25 | | | 22 |
| 18 | 17 [28] | [37] | [35] | 27 | 22 [34] | 39 | 34 |
| 19 | 19 | [37] | [33] | 32 | 46 [36] | 34 | 34 |
| 20 | | [38] | [26] | 31 | 44 [30] | 56 | 15 |
| 22 | 40 [38] | | [42] | 29 | | | 36 |
| 23 | 9 [35] | | [34] | 30 | | | |
| 24 | 38 [32] | | [28] | 26 | | | |
| 25 | 14 [36] | | | | | | |
| 26 | | | [34] | 33 | 48 [36] | 31 | 65 |
| 28 | | | | | 29 | | |
| 29 | | | | | 28 | | |
| 30 | | | | | 38 | | |
| 31 | | | | | 26 | | |
| 32 | | | | | 28 | | |
| 39 | | | | | | 25 | 30 |

**Table S2: Sample sizes for the lab dataset, by population (4 locations) and experimental food treatment (high and low food).** The founders of the lab populations were sampled from either high predation-risk (HP) or low predation-risk (LP) populations, from two distinct river drainage systems (the Oropuche and the Yarra). In total, the lab dataset includes 4629 sets of observations of 211 individuals. Data are from Reznick *et al.* (2004, 2005)

| Population | Food treatment | Number of individuals | Total observations |
|---|---|---|---|
| Oropuche HP | High | 22 | 620 |
| | Low | 26 | 718 |
| Oropuche LP | High | 27 | 450 |
| | Low | 30 | 599 |
| Yarra HP | High | 24 | 615 |
| | Low | 29 | 726 |
| Yarra LP | High | 26 | 428 |
| | Low | 27 | 473 |

### Supporting Figure

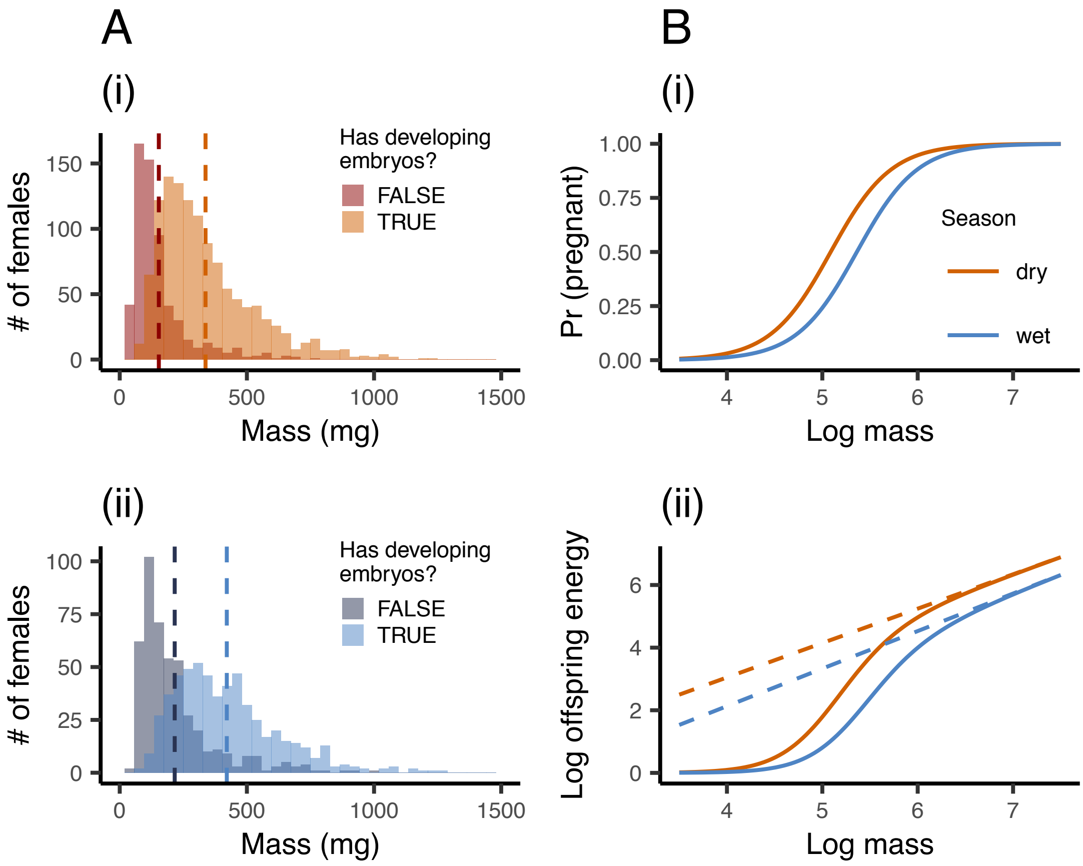


**Figure S1. In wild populations, the probability of an individual being pregnant depended on size and season.** (A) Histograms showing the size distribution of sampled females that contained (light) or did not contain developing embryos (dark) in (i) the dry and (ii) the wet season. Dashed lines indicate distribution means. (B.i) Probability of pregnancy as a function of female size (natural log of mass) in the dry season (red) and the wet season (blue). (B.ii) Log reproduction as a function of log size, when incorporating the probability of pregnancy by size (solid lines) in the dry (red) and wet season (blue). Dashed lines show the relationship between ln(size) and ln(reproduction) given that an individual is pregnant. Solid lines were calculated by multiplying the pregnancy probability function (B.i) by the season-specific log-linear function (dashed lines).
